# The nuclei of human adult stem cells can move within the cell and generate cellular protrusions to contact other cells

**DOI:** 10.1101/2023.07.27.550790

**Authors:** Carlos Bueno, David García-Bernal, Salvador Martínez, Miguel Blanquer, José M. Moraleda

## Abstract

Despite a considerable interest in understanding the mechanisms regulating nucleus structure, chromatin organization and nuclear positioning over decades, the exact significance of the variety of morphologies and positioning that cell nuclei can adopt and their relationship in cellular function is still far from being clearly understood. In this study, we examined the functional significance of the variety of morphologies and positioning that cell nuclei of human bone marrow-derived mesenchymal stem cells can adopt during neural-like differentiation. Here, we show that after neural induction, human bone marrow-derived mesenchymal stem cells enter an intermediate cellular state in which the nuclei are observed to be able to move within the cells, switching shapes and positioning and even generating cellular protrusions as they attempt to contact the cells around them. These findings suggest that changes in nuclear positioning are due to the fact that human cell nuclei somehow sensing their environment.

## Introduction

Since the first observation of a nucleus in 1700^1^, our knowledge of nuclear composition, organization, and positioning has continuously evolved^2,3,4^. Most textbooks depict the nucleus as a spherical or ovoid object at the center of the cell. However, different cell types have very different nuclear shapes and the position of nuclei varies dramatically from this simple view^2,3,5^. Although the cell nucleus has always been considered as the largest and most rigid organelle of eukaryotic cells, emerging views of the nucleus indicate a more dynamic organelle than expected^6^.

There is increasing evidence that nuclei are frequently asymmetrically positioned depending on cell type, developmental stage, migratory state, and differentiation status^2,7,8,9,10^. It has been reported that the position of the nucleus contributes to cell mechanics, as gene regulation through relative genome segregation and the organization of cells within tissues^2,9,10,11^. Furthermore, it is important to note that changes in nuclear morphology and positioning are often associated with cellular dysfunction and disease^2,12,13^. Therefore, the nucleus must be considered not only as the primary site for the storage of genetic material and gene transcription but also as a fundamental mechanical component of the cellular structure^6^. Despite these advances, the exact significance of nuclear positioning in cellular function and tissue physiology is still far from being clearly understood^2,5,7^.

In our laboratory we focus on the differentiation of human mesenchymal stem cells (hMSCs) to generate a neuronal lineage^15,16,17,18^. hMSCs are considered as promising candidates for cell-based regenerative medicine due to their self-renewal capacity, multilineage differentiation potential, trophic effects and immunomodulatory properties^19,20^. Controlled neural differentiation of hMSCs could therefore become an important source of cells for cell therapy of neurodegenerative diseases, as autologous adult hMSCs are easily harvested and effectively expanded^21,22,23^.

Over the past two decades, it has been reported that bone marrow-derived cells (BMDCs) and hMSCs can be induced to overcome their mesenchymal fate and differentiate into neural cells, both *in vitro*^24,25,26,27,28,29,30,31,32^ and *in vivo*^33,34,35,36,37,38,39^, a phenomenon known as transdifferentiation. The term transdifferentiation refers to the conversion of one mature cell type into another cell of different blastodermic origin^40,41^. Such interconversions may involve to regress into an intermediate step before cells differentiate into a new blastodermic potentiality and mature phenotype, or they may occur directly in a process that bypasses such intermediate phenotypes^40,41^. The actual occurrence of neuronal transdifferentiation of BMDCs and MSCs is currently much debated because the findings and their interpretation have been questioned^42,43^. The main argument against these observations in culture studies is that MSCs rapidly adopt neural-like morphologies by retraction of the cytoplasm, rather than by active neurite extension^42,44,45,46^. The *in vivo* neural transdifferentiation of BMDCs and hMSCs has also been questioned, as cell fusion could explain the development of new cell types, that are misinterpreted as transdifferentiated cells^43^.

In a previous publication^18^, we showed that when human bone marrow-derived MSCs (hBM-MSCs) were exposed to a neural induction medium, they rapidly reshaped from a flat to a spherical morphology. Subsequently, hBM-MSCs could maintain the spherical morphology or adopt a new one; they gradually adopted a neural-like morphology through active neurite extension or re-differentiated back to the mesenchymal fate. Furthermore, we found that hBM-MSCs can rapidly and repeatedly switch lineage without cell division. Our results provide evidence that differentiation of hBM-MSCs into neural-like cells requires a transition through a transient and characterized intermediate state of hBM-MSCs (hBM-MSC-derived intermediate cells) and provide a stronger basis for rejecting the idea that the rapid acquisition of a neural-like morphology during MSC transdifferentiation is merely an artifact.

This previous work also highlights that nuclear remodeling occurs during *in vitro* neural-like differentiation of hBM-MSCs. We found that nuclei in hBM-MSC-derived intermediate cells moved within the cell, adopting different morphologies, and even forming two nuclei connected by an internuclear bridge. These nuclear movements generated cellular protrusions that appeared and disappeared from the surface of hBM-MSC-derived intermediate cells. The hBM-MSC- derived intermediate cells positioned their nucleus at the front of the cell during migration. Our results showed that binucleated hBM-MSCs can be formed during neural-like differentiation with independence of any cell fusion, providing evidence that transdifferentiation may also be the mechanism behind the presence of gene-marked binucleated neurons after gene-marked bone marrow-derived cell transplantation. It is important to note that binucleated and polymorphic nuclear cells have been detected in adult neurogenic niches^15,47,48,49,50^.

Taken together, these findings suggest that to date there is no conclusive evidence to continue to consider neuronal transdifferentiation of BMDCs and MSCs as a simple experimental artifact, since it recapitulates some structural steps described *in vivo*^40,41^. Therefore, future studies are needed to understand the mechanisms of these cell conversion processes and eventually harness them for use in regenerative medicine.

In the present study, we investigated the sequence of biological events during neural-like differentiation of hBM-MSCs using live-cell nuclear fluorescence labelling and time-lapse microscopy to understand why the nuclei of hBM-MSC-derived intermediate cells move within the cell generating the cellular protrusions that appear and disappear from the surface.

## Results

### Characterization of hBM-MSCs cultures

In a previous publication^18^, we showed that hBM-MSCs did not express hematopoietic lineage markers such as CD45, CD14, CD34 and CD20, and were positive for CD90, CD105, CD73, thereby demonstrating a characteristic immunophenotype of hMSCs. Under proliferation conditions, hBM-MSCs displayed a flat, fibroblast-like morphology with β-III-tubulin microtubules and actin microfilaments oriented parallel to the longitudinal axis of the cell (Fig. 1a). During interphase, hBM-MSCs displayed a flattened, ellipsoidal nucleus, often located in the center of the cell and with a nuclear volume of approximately 419′30 ± 106′38 μm^3^ (Fig. 1b). The dynamic localization of the nuclear lamina was analyzed by immunostaining for lamin A/C, a nuclear lamina component^51^ and the dynamic localization of the nucleoli was analyzed by immunostaining for fibrillarin, the main component of the active transcription centers^52^. A speckled pattern was observed distributed throughout the nucleus with heterogeneity in the number, size, and distribution of the fibrillarin positive specks (Fig. 1c). Laser scanning confocal microscopy revealed that the inner surface of the nuclear envelope is lined by the nuclear lamina (Fig. 1d).

**Fig. 1.**
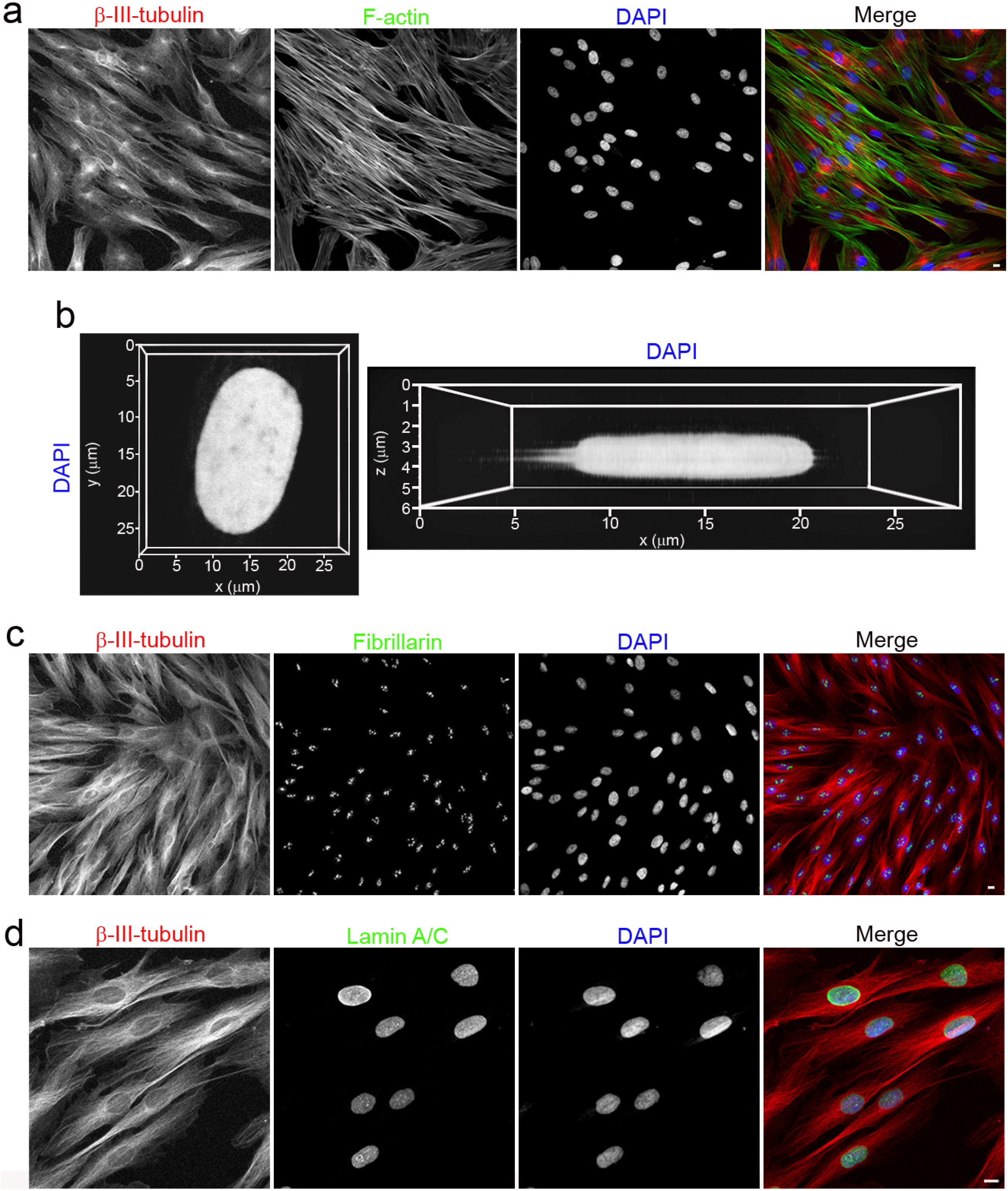
Morphology of hBM-MSCs cultured in basal medium. **a**, Undifferentiated hBM-MSCs exhibited a fibroblast-like morphology with β-III tubulin microtubules and actin microfilaments oriented parallel to the longitudinal axis of the cell. **b**, During interphase, hBM-MSCs displayed a flattened, ellipsoidal nucleus, often located in the center of the cell. **c**, Distribution of fibrillarin-positive specks in the nuclei of undifferentiated hBM-MSCs. **d**, Immunocytochemical analysis revealed that the inner surface of the nuclear envelope is lined by the nuclear lamina. Scale bar: 10 μm.

### Characterization of hBM-MSC-derived intermediate cells

In a previous publication^18^, we showed that following neural induction, hBM-MSCs rapidly reshaped from a flat to a spherical morphology (hBM-MSCs intermediate cells). Subsequently, we observed that hBM-MSC-derived intermediate cells can preserve their spherical shape, change to that of neural-like cells through active neurite extension or revert back to their mesenchymal morphology.

In this study, we focus on hBM-MSC-derived intermediate cells that can maintain their spherical shape for several days without assuming new fates. To better understand why the nuclei of hBM-MSC-derived intermediate cells move within the cell to generate the cellular protrusions, we performed time-lapse microscopy and immunocytochemical analyses of histone H2B-GFP transfected hBM-MSCs within the first 70 hours of neural induction. The time-lapse experiments were performed using a higher magnification objective and a shorter image capture interval than previously published experiments^18^.

Time-lapse imaging revealed that when hBM-MSCs were exposed to a neural induction medium, they rapidly reshaped from a flat to a spherical morphology (Fig. 2a and Supplementary Video 1). We then observed hBM-MSC-derived intermediate cells in which nuclear movements generated only one cell protrusion (Fig. 2a, white arrow and Supplementary Video 1) and hBM-MSC-derived intermediate cells in which nuclear movements alternately generated one or two cellular protrusions (Fig. 2a, yellow arrows and Supplementary Video 1). We found that when hBM-MSC-derived intermediate cell has a nucleus without lobes, its movement within the cell generates only one cell protrusion (Fig. 2b and Supplementary Video 2). However, if the hBM-MSC-derived intermediate cell has a lobed nucleus it will generate one or two cellular protrusions depending on how it moves within the cell (Fig. 2c).

**Fig. 2.**
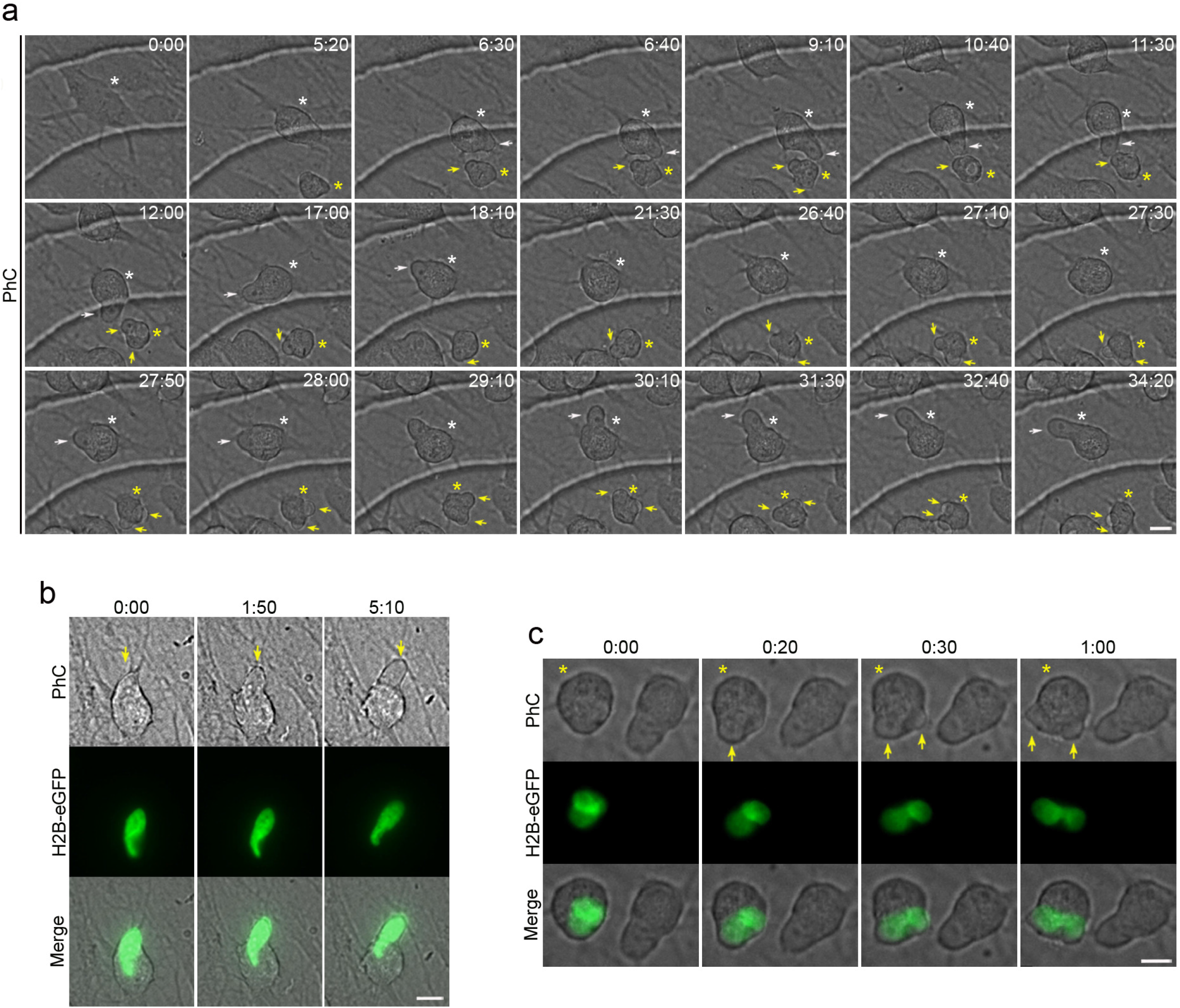
Nuclear movement generated cellular protrusions that appeared and disappeared from the surface of hBM-MSC-derived intermediate cells. **a**, Time-lapse imaging showed that when hBM-MSCs were exposed to a neural induction medium, they rapidly reshaped from a flat to a spherical morphology. Subsequently, we observed hBM-MSC-derived intermediate cells (white asterisk) in which nuclear movements generate only one cell protrusion (white arrow), and hBM-MSC-derived intermediate cells (yellow asterisk) in which nuclear movements alternately generate one or two cellular protrusions (yellow arrows). **b**, We found that when a hBM-MSC-derived intermediate cell has a nucleus without lobes, its movement within the cell generates only one cell protrusion (yellow arrow). **c**, However, if the hBM-MSC-derived intermediate cell has a lobed nucleus it will generate one or two cellular protrusions depending on how it moves within the cell (yellow arrows). Scale bar: 10 μm. PhC: Phase-contrast photomicrographs.

Although the cell nuclei switch their morphologies while moving, time-lapse imaging and immunocytochemical analysis revealed that hBM-MSC-derived intermediate cells have three main different nuclear morphologies: Tail-less nuclei (Fig. 3a, white asterisk), tailed nuclei (Fig. 3a, green asterisk) and lobed nuclei (Fig. 3a, yellow asterisk). Tail-less and tailed nuclei movements generate only a single cell protrusion (Fig. 3a, white and green arrows respectively). However, as mentioned above, lobed nucleus movements generate one or two cellular protrusions depending on how it moves within the cell (Fig. 3a, yellow arrows).

**Fig. 3.**
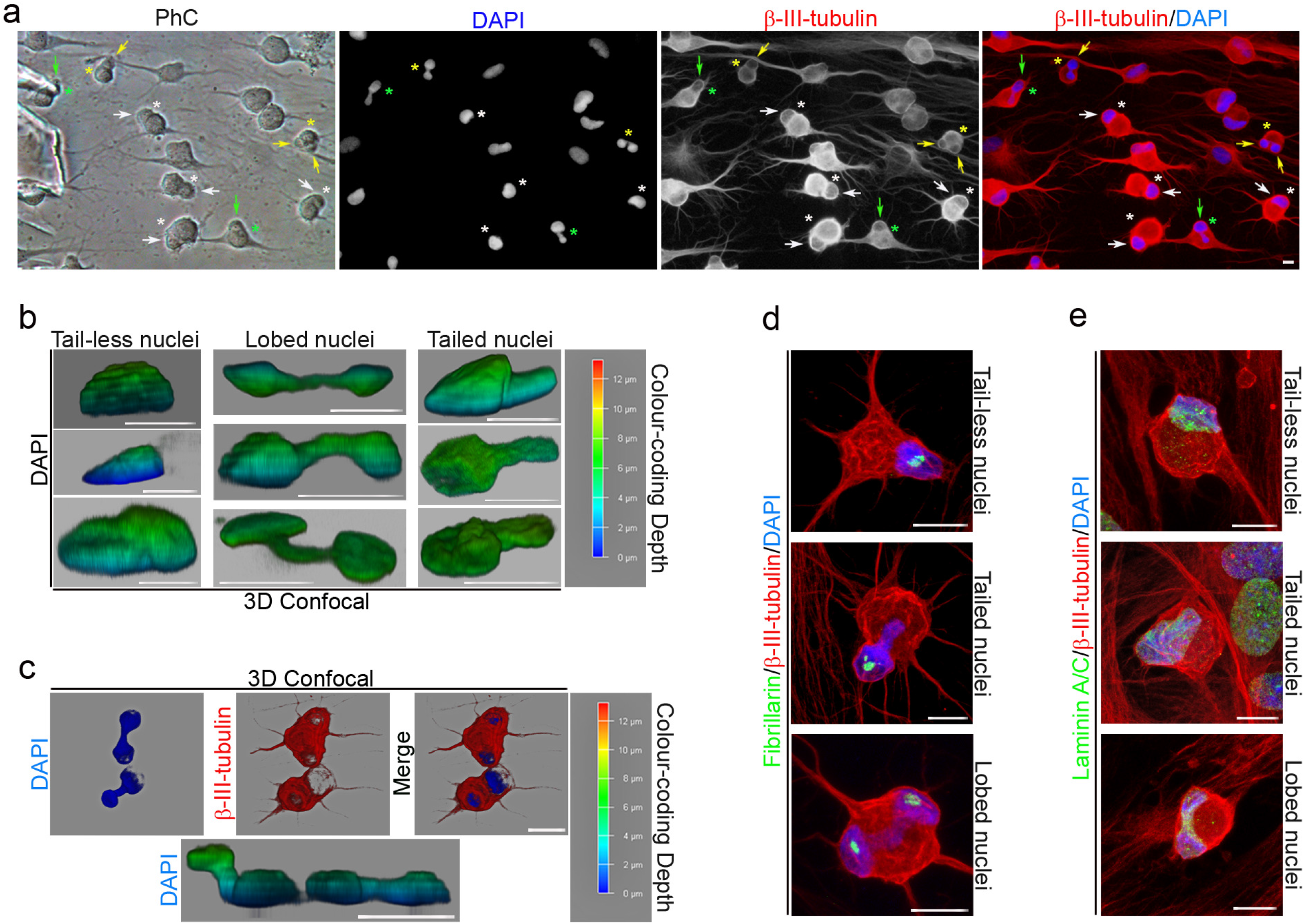
Characterization of hBM-MSC-derived intermediate cell nuclei. **a,** Immunocytochemical analysis revealed that hBM-MSC-derived intermediate cells primarily have three different nuclear morphologies: Tail-less nuclei (white asterisk), tailed nuclei (green asterisk) and lobed nuclei (yellow asterisk). Tail-less and tailed nuclei movements generate only one cell protrusion (white and green arrows respectively). However, lobed nuclei movements generate one or two cellular protrusions depending on how it moves within the cell (yellow arrows). Confocal microscopy analysis and 3D reconstruction revealed that there are small variations in both shape and size within the three types of nuclear morphology (**b**) and that the lobes of the lobed nuclei can be located in different z-planes (**c**). **d**, Immunocytochemical analysis revealed that tail-less nuclei and tailed nuclei contained one or two fibrillarin positive specks while lobed nuclei contained one or two fibrillarin-positive specks in each lobe. **e**, The inner surface of the nuclear envelope is lined by the nuclear lamina. Scale bar: 10 μm. PhC: Phase-contrast photomicrographs.

Confocal microscopy analysis and 3D reconstruction revealed that there are small variations in both shape and size within the three types of nuclear morphology (Fig. 3b). Tail-less nuclei, tailed nuclei and lobed nuclei have a volume of approximately 327′11 ± 94′19 μm^3^, 306′89 ± 16′50 μm^3^ and 361′75 ± 147′44 μm^3^ respectively. It is important to note that confocal microscopic analysis and 3D reconstruction also revealed that the lobes of the lobed nuclei can be located in different z-planes (Fig. 3c).

We observed that tail-less nuclei and tailed nuclei contained one or two fibrillarin positive specks, whereas lobed nuclei contained one or two fibrillarin positive specks in each lobe (Fig. 3d). No positive fibrillarin specks have been detected either in the tail of the tailed nuclei or in the region of nucleoplasm connecting each lobe of lobed nuclei. Laser scanning confocal microscopy also revealed that the inner surface of the nuclear envelope is lined by the nuclear lamina (Fig. 3e).

Confocal microscopy analysis of the cytoskeletal organization of hBM-MSC-derived intermediate cells showed that the cytoskeletal network is reorganized. Actin microfilaments and β-III tubulin microtubules are no longer oriented parallel to the longitudinal axis of the hBM-MSC-derived intermediate cells (Fig. 4a). Furthermore, confocal microscopy analysis and 3D reconstruction revealed that cell protrusions are almost devoid of actin microfilaments and that the β-III tubulin protein is concentrated at the cell protrusion rim (Fig. 4b, black arrows). Although, it has been reported that direct connections between the actin cytoskeleton and the nucleus govern nuclear positioning and nuclear movement during cell polarization and migration^2,7,8^, we observed that the absence of the cytoskeletal network could facilitate nuclear movement within the cell. Future research is required to determine the feasibility of this conjecture.

**Fig. 4.**
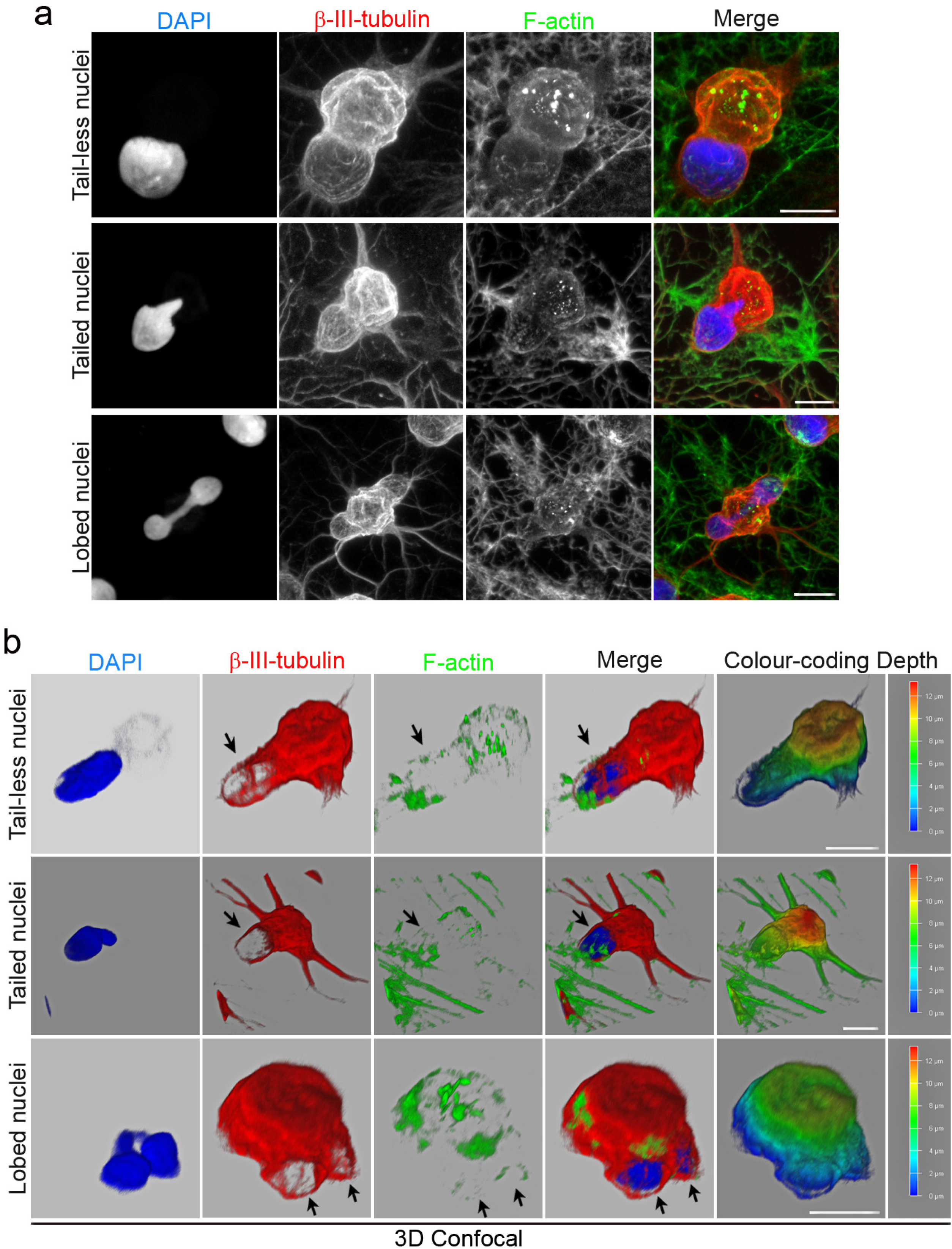
Cytoskeletal organization of hBM-MSC-derived intermediate cells. **a**, Immunocytochemical analysis revealed that actin microfilaments and β-III tubulin microtubules are no longer oriented parallel to the longitudinal axis of the hBM-MSC-derived intermediate cells. **b**. Confocal microscopy analysis and 3D reconstruction revealed that cell protrusions are almost devoid of actin microfilaments and that the β-III tubulin protein is concentrated at the cell protrusion rim (black arrows). Scale bar: 10 μm.

### Nuclear remodeling

To further understand how the three different types of nuclei that move within the hBM-MSC- derived intermediate cells are formed, we examined the sequence of biological events during neural-like differentiation of histone H2B-GFP-transfected hBM-MSCs using time-lapse microscopy. Time-lapse imaging revealed that there are variations in the form and time at which cell nuclei adopt these different nuclear morphologies (Supplementary Videos 3-11). Although future studies are needed to determine all the possible different nuclear remodeling sequences that occur when hBM-MSCs reshape from a flat to a spherical morphology, below we have described some examples of the formation of each of the three types of nuclei that move within hBM-MSC- derived intermediate cells.

We observed that tail-less nuclei are formed by a single nuclear remodeling that occurs as the cell reshapes from a flat to a spherical morphology, positioning the nucleus in a peripheral position within the cell (Fig. 5, white arrows and Supplementary Videos 3,4). The duration of this particular process is approximately 30 hours. Subsequently, the tail-less nucleus began to move within the hBM-MSC-derived intermediate cells. (Fig. 5, yellow arrows and Supplementary Videos 3,4).

**Fig. 5.**
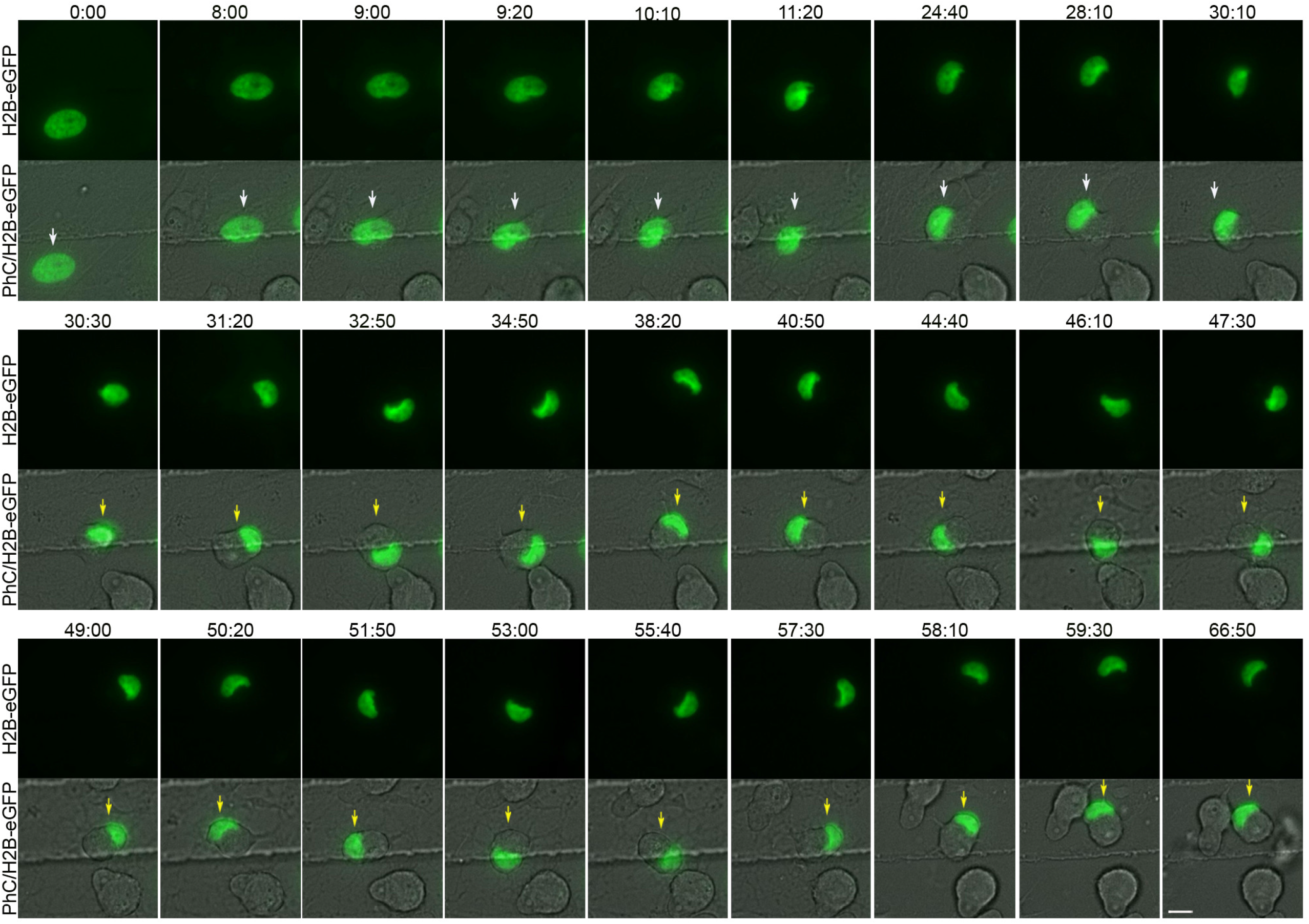
Tail-less nuclei formation. Time-lapse imaging revealed that tail-less nuclei are formed by a single nuclear remodeling that occurs as the cell reshapes from a flat to a spherical morphology, positioning the nucleus in a peripheral position within the cell (white arrows). Subsequently, the tail-less nucleus began to move within the hBM-MSC-derived intermediate cells (yellow arrows). Scale bar: 10 μm.

Time-lapse images also showed that tailed nuclei are formed by one (Supplementary Video 5) or two nuclear remodeling sequences. Below, we show an example of the formation of a tailed nucleus generated by two nuclear remodeling sequences (Fig. 6 and Supplementary Videos 6,7). We found that first a nuclear remodeling sequence occurs as the cell reshapes from a flat to a spherical morphology, positioning the nucleus in a peripheral position within the cell (Fig. 6, white arrows and Supplementary Videos 6,7). The duration of this particular process is approximately 12 hours. Subsequently, the cell nucleus moves (Fig. 6, green arrows and Supplementary Videos 6,7) and undergoes a second nuclear remodeling sequence in which a tailed process in the nucleus is formed (Fig. 6, yellow arrows and Supplementary Videos 6,7). The duration of this particular process is approximately 5 hours. Finally, the tailed nucleus began to move within the hBM-MSC-derived intermediate cells. (Fig. 6, green arrows and Supplementary Videos 6,7). Time-lapse imaging also revealed that as the nuclei move within the cell, the tails can switch shape and size (Fig. 7a and Supplementary Video 8) and even appear to move in different z-planes (Fig. 7b and Supplementary Videos 6,7). Future analyses will be needed to determine whether the nuclei use the tails to stabilize their position within the cell.

**Fig. 6.**
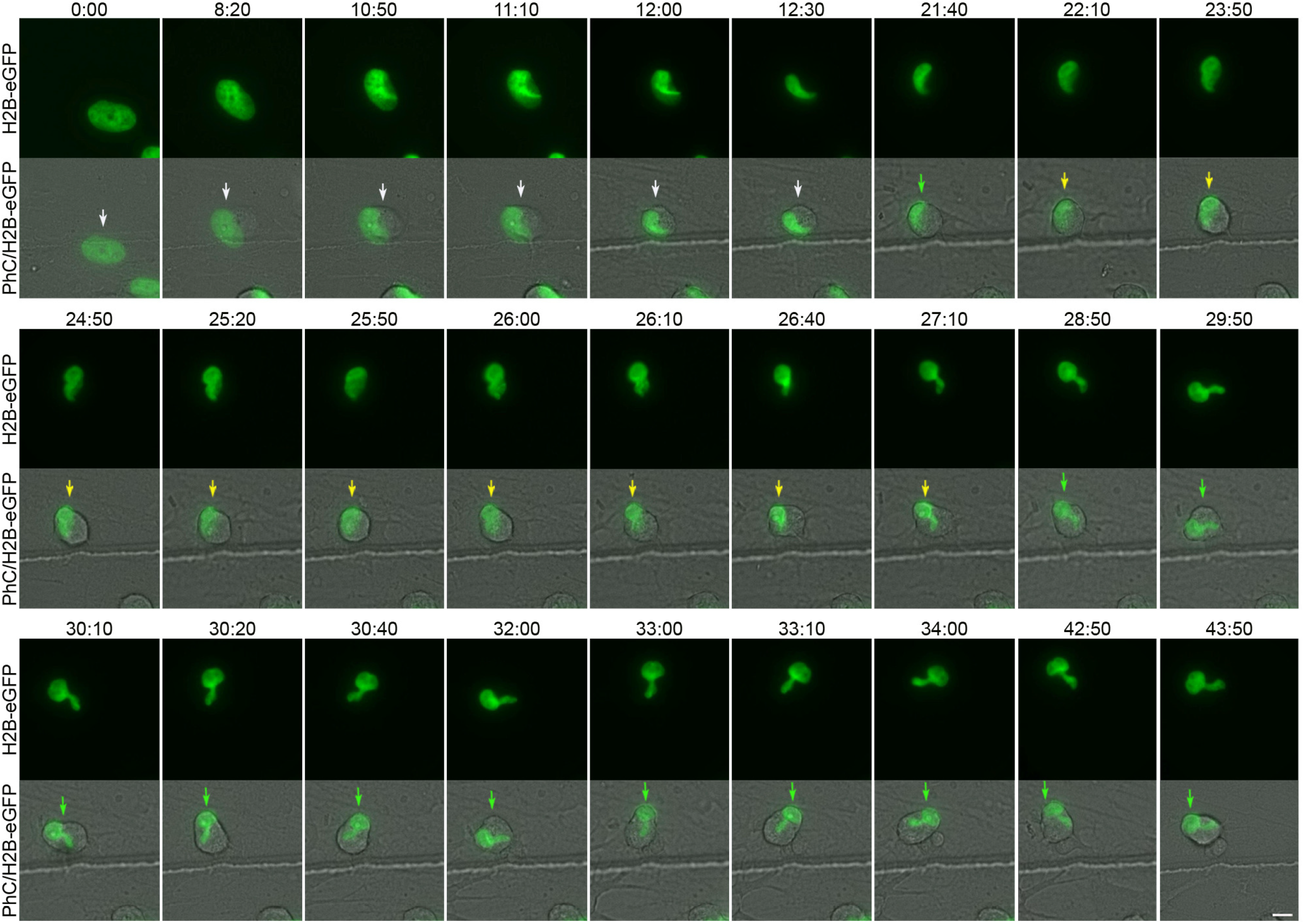
Tailed nuclei formation by two nuclear remodeling sequences. Time-lapse images show that first a nuclear remodeling sequence occurs as the cell reshapes from a flat to a spherical morphology, positioning the nucleus in a peripheral position within the cell (white arrows). Next, the cell nucleus moves (green arrows) and undergoes a second nuclear sequence in which a tailed process in the nucleus is formed (yellow arrows). Finally, the tailed nucleus began to move within the hBM-MSC-derived intermediate cells (green arrows). Scale bar: 10 μm.

**Fig. 7.**
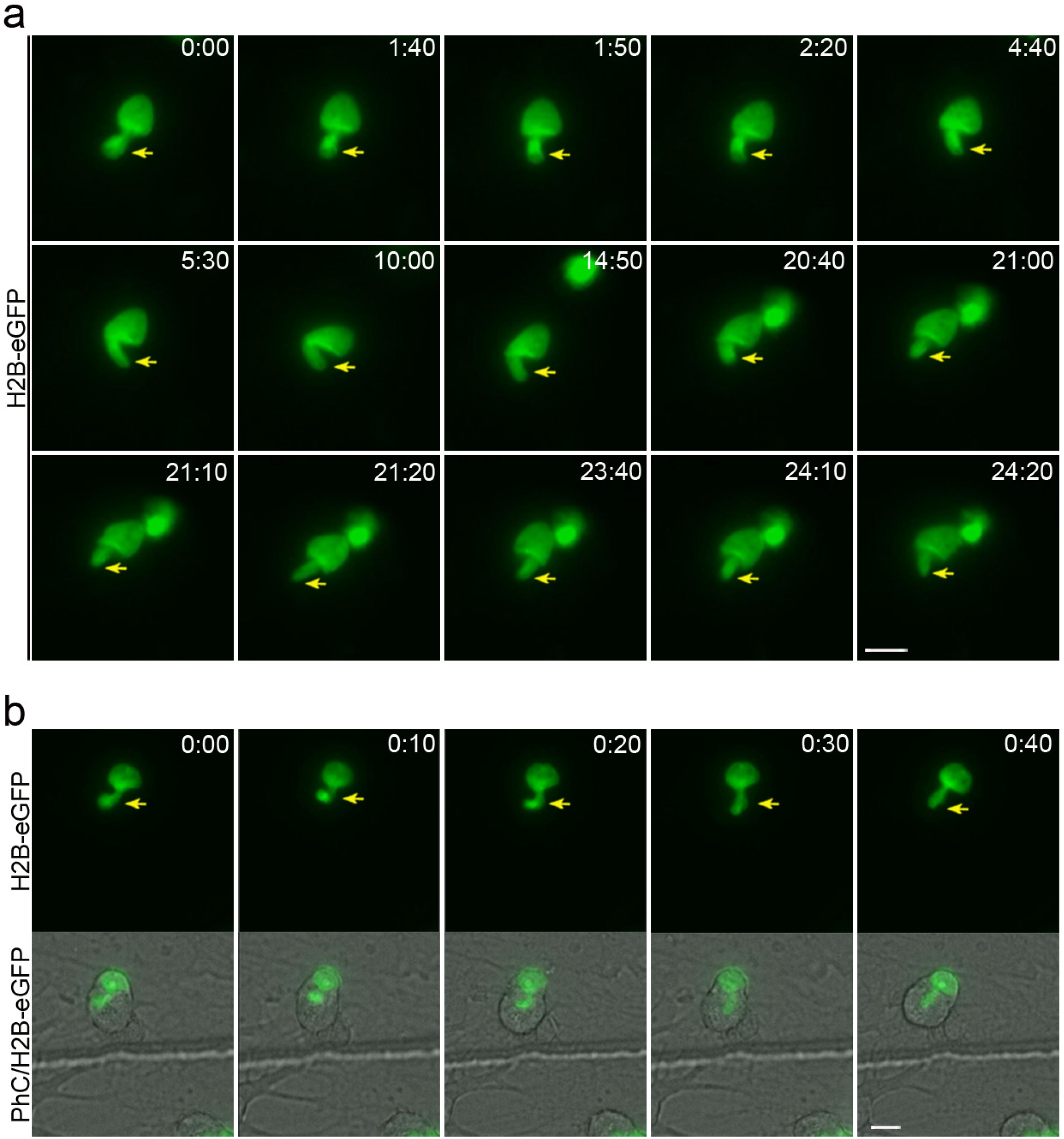
The tails of the tailed nuclei can move within the hBM-MSC-derived intermediate cells, switching shape and size and even move in different z-planes. **a**, Time-lapse imaging revealed that as the tailed nuclei move within the cell, the tails can switch shape and size (yellow arrows). **b**, Time-lapse images also showed that the tails even appear to move in different z-planes. Scale bar: 10 μm.

Time-lapse imaging also revealed that lobed nuclei are formed by one (Supplementary Video 8) or two nuclear remodeling sequences. Below, we show you an example of the formation of a lobed nucleus generated by two nuclear remodeling sequences (Fig. 8 and Supplementary Videos 9,10). We found that first a nuclear remodeling sequence occurs as the cell reshapes from a flat to a spherical morphology, positioning the nucleus in a peripheral position within the cell (Fig. 8, white arrows and Supplementary Videos 9,10). The duration of this particular process is approximately 14 hours. Subsequently, the cell nucleus moves (Fig. 8, green arrows and Supplementary Videos 9,10) and undergoes a second nuclear remodeling sequence in which a lobed nucleus is formed (Fig. 8, yellow arrows and Supplementary Videos 9,10). The duration of this particular process is approximately 5 hours. The lobed nuclei then began to move within the hBM-MSC-derived intermediate cells (Fig. 8, green arrows and Supplementary Videos 9,10). Finally, we also found that lobed nuclei can switch shape while moving within the cell, becoming tailed nuclei (Fig. 8, blue arrows and Supplementary Videos 9,10).

**Fig. 8.**
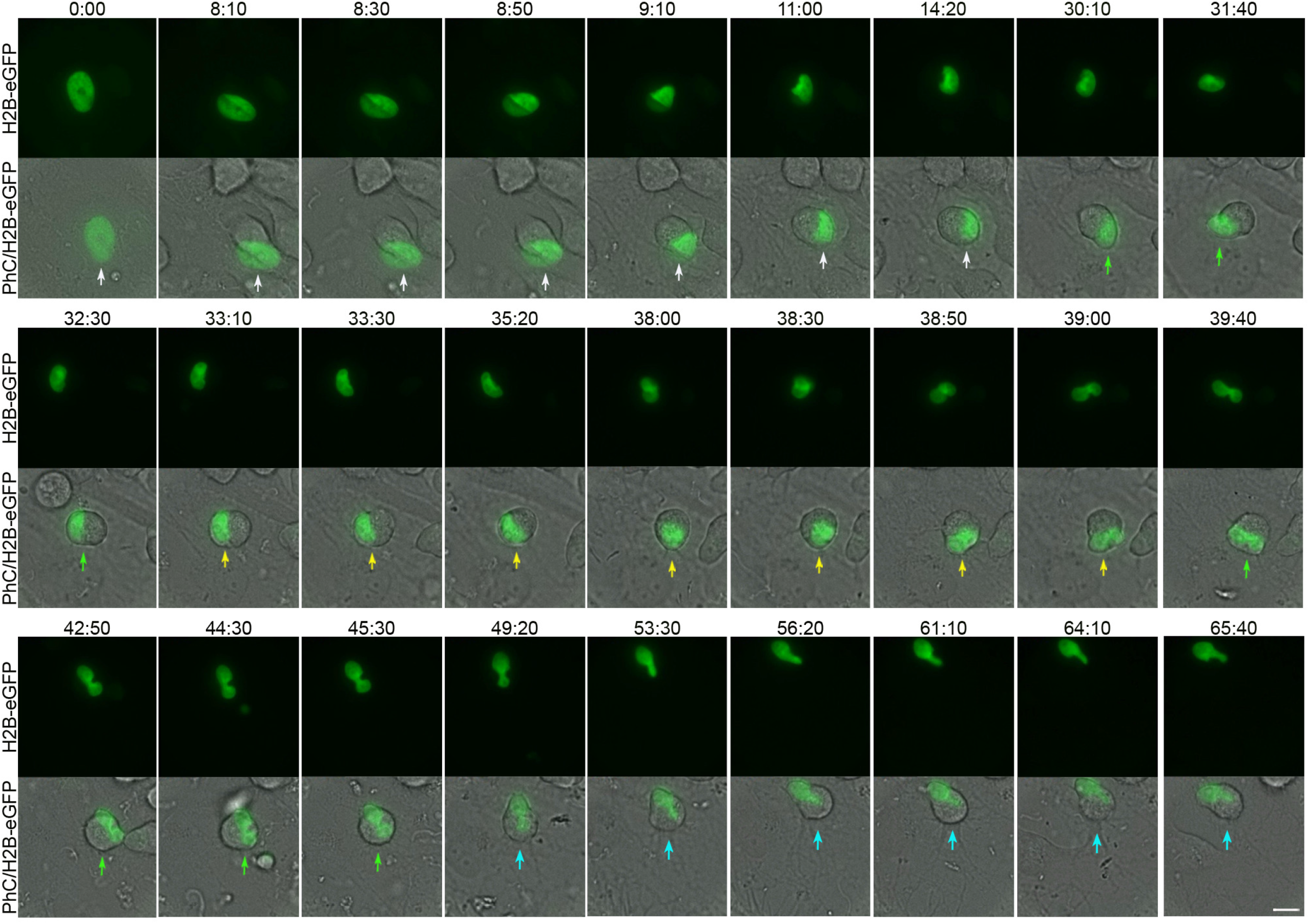
Lobed nuclei formation by two nuclear remodeling sequences. Time-lapse imaging showed that first a nuclear remodeling sequence occurs as the cell reshapes from a flat to a spherical morphology, positioning the nucleus in a peripheral position within the cell (white arrows). Subsequently, the cell nucleus moves (green arrows) and undergoes a second nuclear sequence in which a lobed nucleus is formed (yellow arrows). Afterwards, the lobed nuclei began to move within the hBM-MSC-derived intermediate cells (green arrows). Finally, we also noted that lobed nuclei can switch shape while moving within the cell, becoming tailed nuclei (blue arrows). Scale bar: 10 μm.

As mentioned above, confocal microscopy analysis and 3D reconstruction revealed that the lobes of the lobed nuclei can be located in different z-planes (Fig. 3c). This result together with the time-lapse noted in Figure 2a and Supplementary Video 1 suggest that each lobule of the lobed nuclei can move in different z-planes.

Quantitative analysis of time-lapse imaging revealed that the nuclear speed oscillated between 0.1 and 1.3 μm/min, with an average speed of 0,66 ± 0,45 μm/min. These findings are consistent with previous studies that reported the physical characteristics of typical nuclear movement in different cell types^2^. Nuclear movement can generate cellular protrusions that extend up to a length similar to the cell diameter (Fig. 2,4 and Supplementary Videos 1,2).

### The hBM-MSC-derived intermediate cells use their cell nuclei to interact with other cells

Time-lapse imaging revealed that hBM-MSC-derived intermediate cell nuclei move within the cell, generating cellular protrusions as cell migration breakthrough process to contact surrounding cells (Fig. 2,5,8 and Supplementary Videos 1,2,4,10). Furthermore, we observed that contacts occur between cell nuclei of hBM-MSC-derived intermediate cells, even for several hours (Fig. 9 and Supplementary Video 11). Confocal microscopy analysis (Fig. 10a) and 3D reconstruction (Fig. 10b) revealed that tail-less nuclei, tailed nuclei and lobed nuclei contact each other. We note that lobed nuclei can contact other cells through one lobe or both simultaneously or successively (Fig. 2a and Supplementary Video 1).

**Fig. 9.**
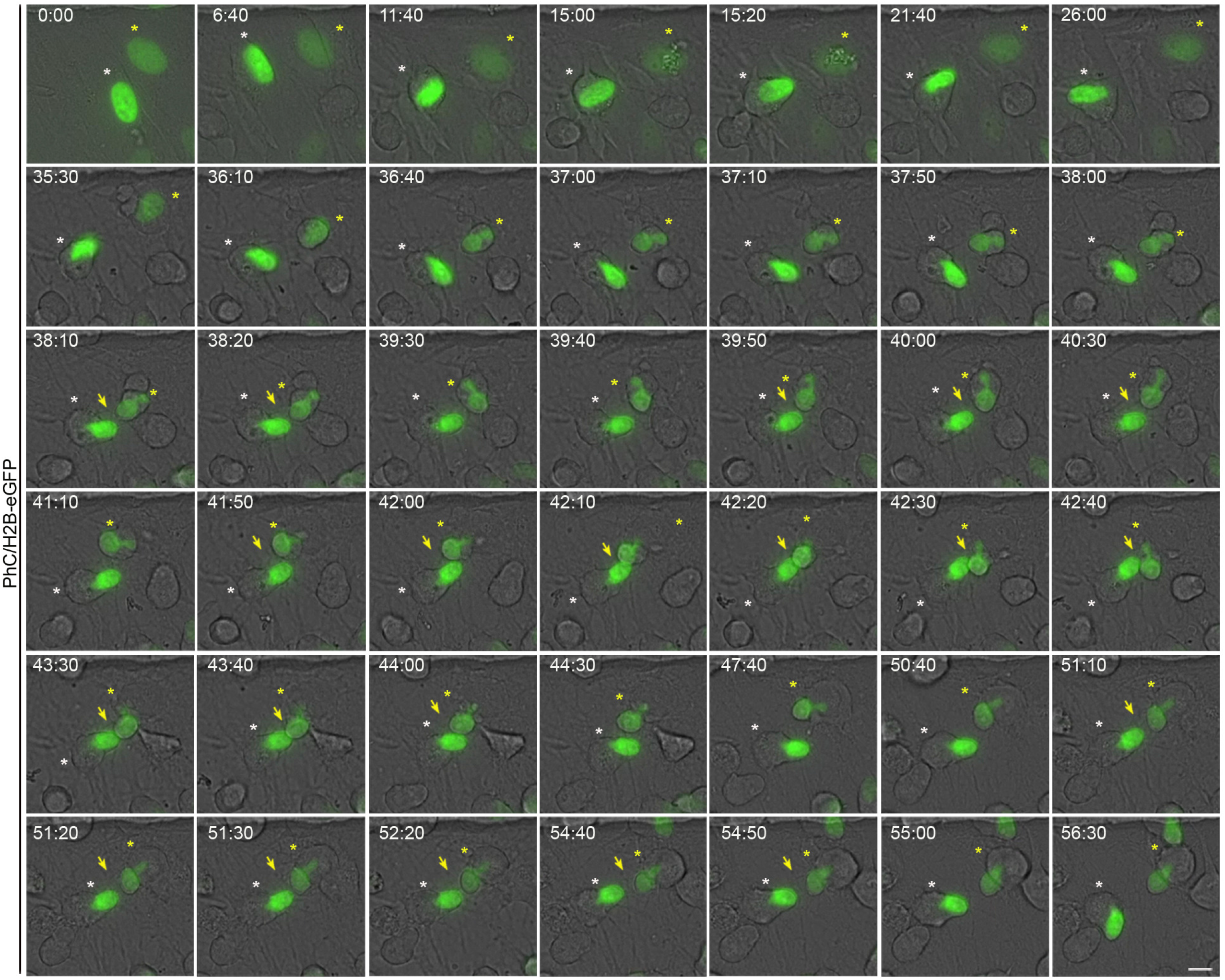
hBM-MSC-derived intermediate cells contact each other through their cell nuclei. Time-lapse imaging revealed that hBM-MSC-derived intermediate cell nuclei move within the cell generating cellular protrusions as they attempt to contact the cells around them, mainly observing contacts between the nuclei (green arrows). Scale bar: 10 μm.

**Fig. 10.**
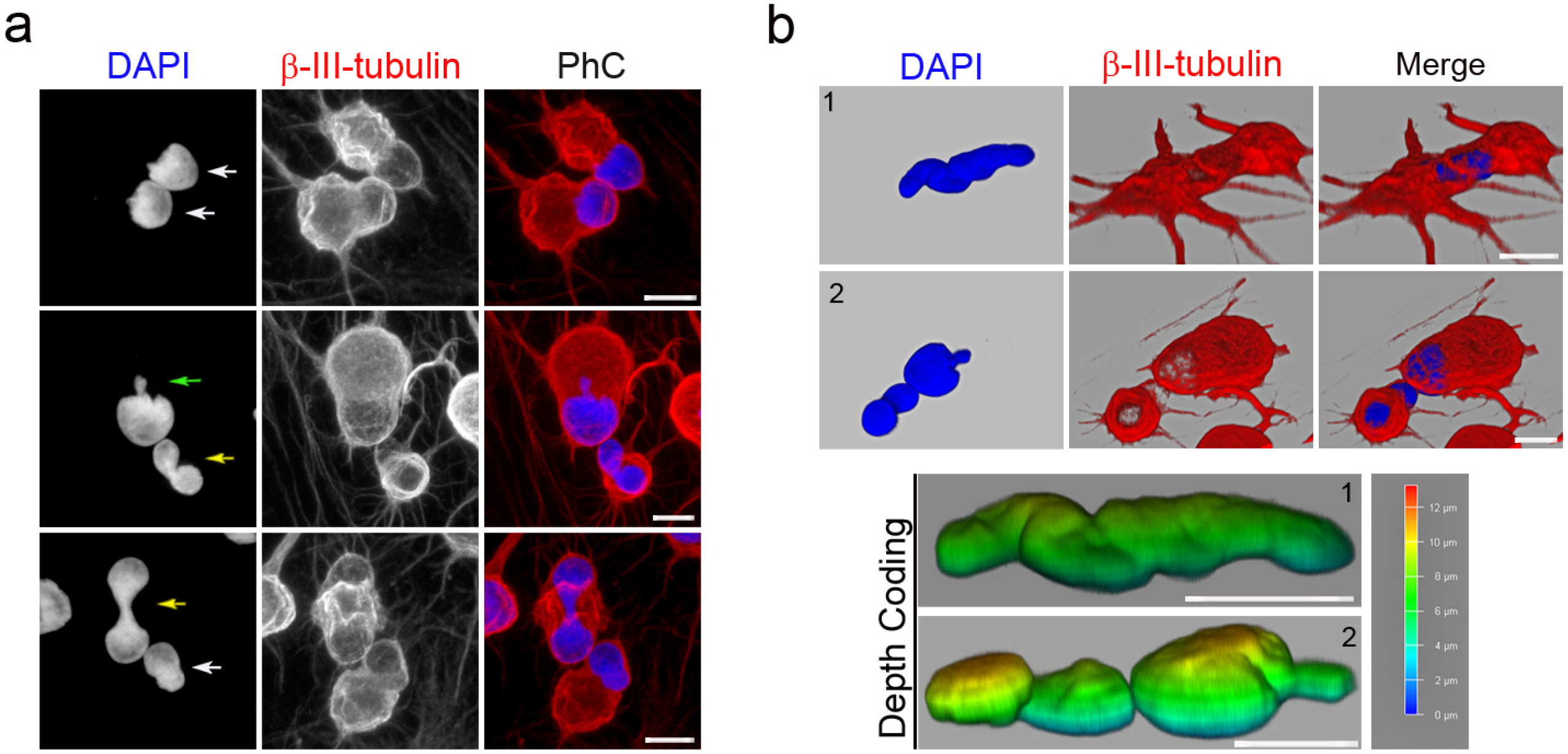
Tail-less nuclei, tailed nuclei and lobed nuclei contact each other. Confocal microscopy analysis (**a**) and 3D reconstruction (**b**) revealed that tail-less nuclei (white arrow), tailed nuclei (green arrow) and lobed nuclei (yellow arrow) contact each other. Scale bar: 10 μm.

Taken together, these findings suggest that changes in nuclear positioning are due to the fact that cell nuclei are somehow sensing their surroundings.

## Discussion

The fate of adult cells was thought to be restricted to their tissue blastodermic of origin^53^. However, there is now a large body of evidence suggesting that under physiological conditions and certain experimental conditions, adult cells may be more plastic than we previously thought in that they can become cells of unrelated lineages, a phenomenon known as transdifferentiation senso stricto^40,41,54,55^. The most evident transdifferentiation events are epitelial-mesenchymal transition (EMT) and its reverse process mesenchymal-epithelial transition (MET)^56,57,58^. These processes are associated with implantation, embryo formation, organ development and wound-healing^56,57,58^ . Importantly, EMT and MET have been implicated in pathological conditions, such as organ fibrosis, and in cancer, where they contributes to tumor progression and metastasis^56,57,58^. In the process of transdifferentiation, cells pass through intermediate states that are not well understood^40,56,57,58,59^. Given the potential application of this cell conversion process, not only in developmental and cancer studies but also in regenerative medicine, a better understanding of intermediate states is crucial to avoid uncontrolled conversion or proliferation, which poses a risk to patients^40,56,57,58,59^.

Over the last two decades, it has been reported that BMDCs and hMSCs can be induced to overcome their mesenchymal fate and transdifferentiate into neural cells, both *in vitro*^24,25,26,27,28,29,30,32,32^ and *in vivo*^33,34,35,36,37,38,39^. However, the neuronal transdifferentiation of BMDCs and MSCs is still considered to be merely an artefact^42,43,46^. The main argument against these observations in culture studies is that MSCs rapidly adopt neuronal-like morphologies by retraction of the cytoplasm, rather than by active neurite extension^42,44,45,46^. The main argument against neuronal transdifferentiation of BMDCs and hMSCs *in vivo* is that cell fusion could explain the development of new cell types, that are misinterpreted as transdifferentiated cells^43^.

In previous publications, we have shown that MSCs isolated from adult human tissues can differentiate into neural-like cells, both *in vitro* and *in vivo*^14,15,16,17,18^. We found that when hBM-MSCs were exposed to a neural induction medium, they rapidly reshaped from a flat to a spherical morphology^18^, Subsequently, hBM-MSCs could maintain the spherical morphology or adopt a new one; they gradually adopted a neural-like morphology through active neurite extension or re-differentiated back to the mesenchymal fate. Furthermore, we found that hBM-MSCs can rapidly and repeatedly switch lineage without cell division. These results provide evidence that the differentiation of hBM-MSCs into neural-like cells requires a transition through an intermediate state, as described in the natural transdifferentiation processes^40,41,56,57,58,59^.

This previous work^18^ also highlights that nuclear remodeling occurs during *in vitro* neural-like differentiation of hBM-MSCs. We found that nuclei in hBM-MSC-derived intermediate cells moved within the cell, adopting different morphologies and even forming two nuclei connected by an internuclear bridge, independently of any cell fusion.

These results provide a strong basis for rejecting the idea that the rapid acquisition of a neural-like morphology during *in vitro* MSC transdifferentiation is merely an artefact and also provide evidence that transdifferentiation may also be the mechanism behind the presence of gene-marked binucleated neurons after gene-marked bone marrow-derived cell transplantation.

It is important to note that the studies describing the presence of binucleated Purkinje neurons after bone marrow-derived cell transplantation suggest, but do not conclusively demonstrate, that cell fusion is the underlying mechanism to explain the presence of binucleated neurons^43,60,61^. Futhermore, it has been reported that binucleated Purkinje neurons are also present in healthy, unmanipulated mice and humans^43,62^. In addition, many authors have described binucleated neurons in various central and peripheral parts of the nervous system^63,64,65,66^. What is more, many authors have reported that many cultured hippocampal neurons^48^ (their Fig. S2), and neural stem cells located in the ventricular-subventricular zone of the anterolateral ventricular wall of the human fetal brain^49^ (their Fig. 2C) and adult mouse brain^50,51^ (their Fig. 1E, 4I, 6F, S1B, S2B, S6A and their Fig. 3A respectively) have two nuclei connected by an internuclear bridge.

Taken together, these results suggest that to date there is no conclusive evidence to date to continue to consider the neuronal transdifferentiation of BMDCs and hMSCs as an artefact. Therefore, future studies are needed to optimize the various neural induction protocols that have been developed for MSCs^67^, not only to understand the mechanisms of these cellular conversion processes, but also to eventually harness them for use in regenerative medicine^40,68,69^.

Our current research aims to improve our understanding of what hBM-MSCs look like at this intermediate stage. We wanted to know why the nuclei of hBM-MSC-derived intermediate cells move within the cell and generate the cellular protrusions that appear and disappear from the surface.

In this study, we have shown that once the hBM-MSCs enter this intermediate stage, the cell nuclei begin to move within the cell and generate the cellular protrusions as they try to contact the surrounding cells. Interkinetic nuclear migration (INM) in neural progenitors underlay normal neurogenesis during normal development^70,71^, suggesting that the observed nuclear motility in hBM-MS-derived intermediated cells recapitulates an intermediate stage before to decide proliferative of differentiative progression in development and transdifferentiation. Importantly, in a previous publication^18^ (their Fig. S4 and Supplementary Video 4), we observed that the nuclei of hBM-MSCs can also exhibit oscillatory movement along the axis of cell polarity, similar to that described to occurs during INM.

We also found that contacts occur between the cell nuclei of hBM-MSC-derived intermediate cells, even for several hours. These findings suggest that changes in nuclear positioning occur because cell nuclei somehow sense their surroundings. Under our knowledge, we have described for the first time the process of direct interactions between cells nuclei, which opens the possibility of a new level of intercellular interaction. During this intermediate phase, nuclear factors may be interchanged by neighboring cells and directly modify transcriptomes. Extensive studies are required to determine the mechanisms and consequences of this process of direct interactions between cells nuclei in normal and pathologic circumstances.

This work also highlights that cell nuclei moving within hBM-MSC intermediate cells have mainly three different morphologies: tail-less nuclei, tailed nuclei, and lobed nuclei. We have shown that there are variations in the shape and time at which the nuclei of hBM-MSC adopt the different nuclear morphologies observed in hBM-MSC-derived intermediate cells. We also found that tail-less and tailed nuclei movements generate only a single cell protrusion when attempting to contact other cells. However, lobed nuclei movements generate one or two cellular protrusions depending on how it moves within the cell. Lobed nuclei may contact other cells through one lobe or both simultaneously or successively. Nevertheless, it is important to note that the three different nuclear morphologies observed in hBM-MSC intermediate cells can be interchanged as the nucleus moves within the cell.

Beyond the nervous system, the presence of lobed nuclei has been reported in most immune cells^3,72^ and in various human organs, including the heart and liver^73,74^. However, the functional significance of multilobed nuclear structures is not yet known^3,72,73,74^. It would therefore be interesting to investigate whether these nuclear structures are also associated with nuclear movement within the cell.

In conclusion, our findings enrich the understanding of intermediate states in the neural-like differentiation process of hBM-MSCs and suggest that changes in nuclear positioning are due to the fact that human cell nuclei somehow sensing their environment. Although future experiments will be needed to understand why there are different nuclear morphologies in hBM-MSC-derived intermediate cells and why they are formed in different ways, our results demonstrate that the nucleus should already be considered not only as the primary site for the storage of genetic material and the transcription of genes, but also as a fundamental mechanical component of the cell. Human mesenchymal stromal cells could not only help to increase our understanding of the mechanisms underlying cellular plasticity, and eventually harness them for use in regenerative medicine, but also facilitate an understanding of the mechanisms regulating nuclear structure and dynamics.

## Methods

### Ethical conduct of research

The authors declare that all protocols used to obtain and process all human samples were approved by the local ethics committees of the Miguel Hernández University of Elche (No. UMH.IN.SM.03.16) and University of Murcia (HULP3617.05/07/2012; HUSA19/1531.17/02/2020) according to Spanish and European legislation and conformed to the ethical guidelines of the Helsinki Declaration. Donors provided written informed consent before obtaining samples.

### Isolation and culture of hBM-MSCs

A standard protocol for isolation and expansion of hMB-MSCs was used as previously described^18^. Bone marrow aspirates were obtained by percutaneous direct aspiration from the iliac crest of 5 healthy volunteers at University Hospital Virgen de la Arrixaca (Murcia, Spain). Bone marrow was collected with 20 U/ml sodium heparin, followed by a Ficoll density gradient-based separation by centrifugation at 540g for 20 min. After, mononuclear cell fraction was collected, washed twice with Ca^2+^/Mg^2+^-free phosphate buffered saline (PBS) (Gibco Invitrogen) and seeded into 175-cm2 culture flasks (Nunc, Thermo Fisher Scientific) at a cell density 1.5×10^5^ cells/cm^2^ in serum-containing media (designated as the basal media), composed of DMEM low glucose medium (Thermo Fisher Scientific) supplemented with 10% fetal bovine serum (FBS; Lonza), 1% GlutaMAX (Thermo Fisher Scientific), non-essential amino acid solution (Sigma-Aldrich) and 1% penicillin/streptomycin (Thermo Fisher Scientific). After 3 days of culture at 37°C and 7% CO_2_, non-attached cells were removed and fresh complete medium was added. Culture media were renewed every 2 days, and the isolated hMB-MSCs were passaged when cultures were 70-80% confluent. All studies were performed using hMB-MSCs expanded within culture passages 3-4.

### Expression Vectors and Cell Transfection

The expression vectors used in the present study were H2B-eGFP, a gift from Geoff Wahl (Addgene plasmid # 11680; http://n2t.net/addgene:11680; RRID:Addgene_11680; Kanda et al^75^., 1998). A standard protocol for transfecting MSCs was used as previously described^18^. Isolated hMB-MSCs were transfected using the Gene Pulser-II Electroporation System (Bio-Rad Laboratories). Electroporation was performed in a sterile cuvette with a 0.4-cm electrode gap (Bio-Rad Laboratories), using a single pulse of 270 V, 500 μF. Plasmid DNA (5 μg) was added to 1.5 × 10^6^ viable hMB-MSCs in 0.2-ml DMEM low glucose medium (Thermo Fisher Scientific) before electrical pulsing.

### Time-lapse microscopy of histone H2B-GFP expressing hBM-MSCs cultured in neural induction media

We used μ-Dish 35 mm, high Grid-500 (Ibidi) for live cell imaging. Histone H2B-GFP transfected hBM-MSCs were plated onto collagen IV (Sigma-Aldrich) coated plastic or glass coverslips. To induce neural differentiation, cells at passage 3–4 were allowed to adhere to the plates overnight. Basal media was removed the following day and the cells were cultured for 2 days in serum-free media (designated as the neural basal media) consisting in Dulbecco’s modified Eagle’s medium/F12 (DMEM/F12 Glutamax, Gibco) supplemented with N2-supplement (R&D systems), 0.6% glucose (Sigma-Aldrich), 5mM HEPES (Sigma-Aldrich), 0.5% human serum albumin (Sigma-Aldrich), 0.0002% heparin (Sigma-Aldrich), non-essential amino acid solution (Sigma-Aldrich) and 100 U/ml penicillin-streptomycin (Sigma-Aldrich). On day 3, cells were cultured in neural induction media, consisting in the neural basal media supplemented with 500nM retinoic acid (Sigma-Aldrich) and 1mM dibutyryl cAMP (Sigma-Aldrich). Time-lapse analysis was carried out using a Widefield Leica Thunder-TIRF imager microscope. We perform time-lapse microscopy within the first 70 hours after neural induction media was added directly to the cells. Time-lapse images were obtained with a 40X objective every 10 min. During imaging, cells were enclosed in a chamber maintained at 37°C under a humidified atmosphere of 5% CO2 in air. Data are representative of ten independent experiments.

### Immunocytochemistry

A standard immunocytochemical protocol was used as previously described^14,15,17,18^. Histone H2B-GFP transfected hBM-MSCs were plated onto collagen IV (Sigma-Aldrich) coated plastic or glass coverslips and maintained in neural induction media. Cells were rinsed with PBS and fixed in freshly prepared 4% paraformaldehyde (PFA; Sigma-Aldrich). Fixed cells were blocked for 2 h in PBS containing 10% normal horse serum (Gibco) and 0.25% Triton X-100 (Sigma) and incubated overnight at 4 °C with antibodies against β-III-tubulin (TUJ1; 1:500, Covance), Fibrillarin (1/300, Abcam) and Lamin A/C (1/300, GeneTex), in PBS containing 1% normal horse serum and 0.25% Triton X-100. On the next day, cells were rinsed and incubated with the secondary antibodies conjugated with Alexa Fluor® 488 (anti-rabbit; 1:500, Molecular Probes) and Alexa Fluor® 594 (anti-mouse; 1:500, Molecular Probes). Cell nuclei were counterstained with DAPI (0.2 mg/ml in PBS, Molecular Probes). Alexa Fluor 488® phalloidin (Molecular Probes) was used to selectively stains F-actin. Data are representative of ten independent experiments per condition.

### Images and Data Analyses

Photograph of visible and fluorescent stained samples were carried out in a Widefield Leica Thunder-TIRF imager microscope equipped with a digital camera or in confocal laser scanning microscope Leica TCS-SP8. We used Leica Application Suite X and Imaris software for image analysis and Filmora Video Editor software for video editing. Photoshop software was used to improve the visibility of fluorescence images without altering the underlying data. Data are representative of ten independent experiments per condition and are expressed as mean ± SD.

## Data availability statement

All data generated or analysed during this study are included in this published article (and its supplementary information files).

## Acknowledgements

This study was supported by the Spanish Ministry of Science and Innovation, the Carlos III Health Institute (ISCIII), through The Spanish Network of Advanced Therapies (TERAV), projects RD21/0017/0001 and RD21/0017/0017, co-funded by European Union “NextGenerationEU. Plan de Recuperación Transformación y Resiliencia”. It was also partially supported by Research Grant of the Universidad Católica San Antonio de Murcia (UCAM). We greatly appreciate the technical assistance of Microscopy Section and the Tissue Culture Service of the University of Murcia.

## Author contributions

C.B. conceived of the study, designed the study, carried out the molecular lab work and drafted the manuscript; D.G-B and M.B. designed experiments, participated in data analysis and helped draft the manuscript; S.M. and J.M.M. helped draft the manuscript and financial support.

## Materials & Correspondence

Carlos Bueno, PhD., Medicine Department and Hematopoietic Transplant and Cellular Therapy Unit, Institute of Biomedical Research (IMIB), University of Murcia, Faculty of Medicine, Murcia, 30120, Spain. Tel.: 0034-968885013. Fax: 00-34-868884150.

## References

1. Osorio, D. S. & Gomes, E. R. The contemporary nucleus: a trip down memory lane. Biol. Cell. 105, 430–441 (2013).

2. Gundersen, G. G. & Worman, H. J. Nuclear positioning. Cell. 152, 1376–1389 (2013).

3. Skinner, B. M. & Johnson, E. E. Nuclear morphologies: their diversity and functional relevance. Chromosoma. 126, 195–212 (2017).

4. Misteli, T. The Self-Organizing Genome: Principles of Genome Architecture and Function. Cell. 183, 28–45 (2020).

5. Jevtić, P, Edens L. J., Vuković, L. D. & Levy, D. L. Sizing and shaping the nucleus: mechanisms and significance. Curr. Opin. Cell Biol. 28, 16–27 (2014).

6. Kalukula Y, Stephens AD, Lammerding J, Gabriele S. Mechanics and functional consequences of nuclear deformations. Nat. Rev. Mol. Cell Biol. 23, 583–602 (2022).

7. Dupin, I. & Etienne-Manneville, S. Nuclear positioning: mechanisms and functions. Int. J. Biochem. Cell Biol. 43, 1698–1707 (2011).

8. Calero-Cuenca, F. J., Janota, C. S. & Gomes, E. R. Dealing with the nucleus during cell migration. Curr. Opin. Cell Biol. 50, 35–41 (2018).

9. Almonacid, M., Terret, M. E. & Verlhac, M. H. Nuclear positioning as an integrator of cell fate. Curr. Opin. Cell Biol. 56, 122–129 (2019).

10. Deshpande, O. & Telley, I. A. Nuclear positioning during development: Pushing, pulling and flowing. Semin. Cell Dev. Biol. 120,10–21 (2021).

11. Bone, C. R. & Starr, D. A. Nuclear migration events throughout development. J. Cell Sci. 129, 1951–1961 (2016).

12. Zwerger, M., Ho, C. Y. & Lammerding, J. Nuclear mechanics in disease. Annu. Rev. Biomed. Eng. 13, 397–428 (2011).

13. Manda, N. K., Golla, U., Sesham, K., Desai, P., Joshi, S., Patel, S., Nalla, S., Kondam, S., Singh, L., Dewansh, D., Manda, H. & Rokana N. Tuning between Nuclear Organization and Functionality in Health and Disease. Cells. 2023 Feb 23;12(5):706. doi: 10.3390/cells12050706.

14. Bueno, C., Ramirez, C., Rodríguez-Lozano, F. J, Tabarés-Seisdedos, R., Rodenas, M., Moraleda, J. M., Jones, J. R. & Martinez, S. Human adult periodontal ligament-derived cells integrate and differentiate after implantation into the adult mammalian brain. Cell Transplant. 22, 2017–2028 (2013).

15. Bueno, C., Martínez-Morga, M. & Martínez, S. Non-proliferative neurogenesis in human periodontal ligament stem cells. Sci. Rep. 2019 Dec 2;9(1):18038. doi: 10.1038/s41598-019-54745-3.

16. Bueno, C. & Martínez, S. Neurogenesis similarities in different human adult stem cells. Neural Regen. Res. 2021 Jan;16(1):123–124. doi: 10.4103/1673-5374.286967.

17. Bueno, C., Martínez-Morga, M., García-Bernal, D., Moraleda, J. M. & Martínez, S. Differentiation of human adult-derived stem cells towards a neural lineage involves a dedifferentiation event prior to differentiation to neural phenotypes. Sci. Rep. 2021 Jun 8;11(1):12034. doi: 10.1038/s41598-021-91566-9.

18. Bueno, C., Blanquer, M., García-Bernal, D., Martínez, S. & Moraleda, J. M. Binucleated human bone marrow-derived mesenchymal cells can be formed during neural-like differentiation with independence of any cell fusion events. Sci. Rep. 2022 Nov 30;12(1):20615. doi: 10.1038/s41598-022-24996-8.

19. Pittenger, M. F., Discher, D. E., Péault, B. M., Phinney, D. G., Hare, J. M. & Caplan, A. I. Mesenchymal stem cell perspective: cell biology to clinical progress. NPJ Regen. Med. 2019 Dec 2;4:22. doi: 10.1038/s41536-019-0083-6.

20. García-Bernal, D. et al. The Current Status of Mesenchymal Stromal Cells: Controversies, Unresolved Issues and Some Promising Solutions to Improve Their Therapeutic Efficacy. Front Cell Dev. Biol. 2021 Mar 16;9:650664. doi: 10.3389/fcell.2021.650664.

21. Mollinari, C., Zhao, J., Lupacchini, L., Garaci, E., Merlo, D. & Pei, G. Transdifferentiation: a new promise for neurodegenerative diseases. Cell Death Dis. 2018 Aug 6;9(8):830. doi: 10.1038/s41419-018-0891-4.

22. Choudhary, P., Gupta, A. & Singh, S. Therapeutic advancement in neuronal transdifferentiation of mesenchymal stromal cells for neurological disorders. J. Mol. Neurosci. 10.1007/s12031-020-01714-5 (2020).

23. Hernández, R. et al. Differentiation of human mesenchymal stem cells towards neuronal lineage: clinical trials in nervous system disorders. Biomol. Ther. 28, 34–44 (2020).

24. Woodbury, D., Schwarz, E. J., Prockop, D. J. & Black I. B. Adult rat and human bone marrow stromal stem cells differentiate into neurons. J. Neurosci. Res. 61, 364–370 (2000).

25. Woodbury, D., Reynolds, K. & Black, I. B. Adult bone marrow stromal stem cells express germline, ectodermal, endodermal, and mesodermal genes prior to neurogenesis. J. Neurosci. Res. 69, 908–917 (2002).

26. Muñoz-Elias, G., Woodbury, D. & Black, I. B. Marrow stromal cells, mitosis and neuronal differentiation: stem cell and precursor functions. Stem Cells. 21,437–448 (2003).

27. Jeong, J. A. et al. Rapid neural differentiation of human cord blood-derived mesenchymal stem cells. Neuroreport. 15, 1731–1734 (2004).

28. Suon, S. et al. Transient differentiation of adult human bone marrow cells into neuron-like cells in culture: development of morphological and biochemical traits is mediated by different molecular mechanisms. Stem Cells Dev. 13, 625–635 (2004).

29. Hermann, A. et al. Comparative analysis of neuroectodermal differentiation capacity of human bone marrow stromal cells using various conversion protocols. J. Neurosci. Res. 83, 1502–1514 (2006).

30. Ning, H., Lin, G., Lue, T. F. & Lin, C. S. Neuron-like differentiation of adipose tissue-derived stromal cells and vascular smooth muscle cells. Differentiation. 74, 510–518 (2006).

31. Azedi, F. et al. Comparative capability of menstrual blood versus bone marrow derived stem cells in neural differentiation. Mol. Biol. Rep. 44, 169–182 (2017).

32. Radhakrishnan, S. et al. In vitro transdifferentiation of human adipose tissue-derived stem cells to neural lineage cells -a stage-specific incidence. Adipocyte. 8, 164–177 (2019).

33. Azizi, A. S., Stokes, D., Augelli, B. J., DiGirolamo, C. & Prockop, D. J. Engraftment and migration of human bone marrow stromal cells implanted in the brains of albino rats-Similarities to astrocyte grafts. Proc. Natl. Acad. Sci. U.S.A. 95, 3908–3913 (1998).

34. Kopen, G. C., Prockop, D. J. & Phinney, D. G. Marrow stromal cells migrate throughout forebrain and cerebellum, and they differentiate into astrocytes after injection into neonatal mouse brains. Proc. Natl. Acad. Sci. U.S.A. 96, 10711–10716 (1999).

35. Brazelton, T. R., Rossi, F. M., Keshet, G. I. & Blau, H. M. From marrow to brain: Expression of neuronal phenotypes in adult mice. Science. 290, 1775–1779 (2000).

36. Mezey, E., Chandross, K. J., Harta, G., Maki, R. A. & McKercher, S. R. Turning blood into brain: Cells bearing neuronal antigens generated in vivo from bone marrow. Science 290, 1779–1782 (2000).

37. Priller, J. et al. Neogenesis of cerebellar Purkinje neurons from gene-marked bone marrow cells in vivo. J. Cell Biol. 155, 733–738 (2001).

38. Mezey, E. et al. Transplanted bone marrow generates new neurons in human brains. Proc. Natl. Acad. Sci. U.S.A. 100, 1364–1369 (2003).

39. Muñoz-Elias, G., Marcus, A. J., Coyne, T. M., Woodbury, D. & Black, I. B. Adult bone marrow stromal cells in the embryonic brain:Engraftment, migration, differentiation, differentiation, and long-term survival. J. Neurosci. 24, 4585–4595 (2004).

40. Jopling, C., Boue, S., & Izpisua Belmonte J. C. Dedifferentiation, transdifferentiation and reprogramming: three routes to regeneration. Nat. Rev. Mol. Cell Biol. 12, 79–89 (2011).

41. Merrell, A. J. & Stanger, B. Z. Adult cell plasticity in vivo: de-differentiation and transdifferentiation are back in style. Nat. Rev. Mol. Cell Biol. 17, 413–25 (2016).

42. Lu, P., Blesch, A. & Tuszynski, M. H. Induction of bone marrow stromal cells to neurons: differentiation, transdifferentiation, or artifact? J. Neurosci. Res. 77, 174–191 (2004).

43. Kemp, K., Wilkins, A. & Scolding, N. Cell fusion in the brain: Two cells forward, one cell back. Acta Neuropathol. 128, 629–638 (2014).

44. Neuhuber, B. et al. Reevaluation of in vitro differentiation protocols for bone marrow stromal cells: Disruption of actin cytoskeleton induces rapid morphological changes and mimics neuronal phenotype. J. Neurosci. Res. 77, 192–204 (2004).

45. Bertani, N., Malatesta, P., Volpi, G., Sonego, P. & Perris, R. Neurogenic potential of human mesenchymal stem cells revisited: Analysis by immunostaining, time-lapse video and microarray. J. Cell Sci. 118, 3925–3936 (2005).

46. Krabbe, C., Zimmer, J. & Meyer, M. Neural transdifferentiation of mesenchymal stem cells-A critical review. APMIS. 113, 831–844 (2005).

47. Doetsch, F., Garcia-verdugo, J. M. & Alvarez-buylla, A. Cellular composition and three-dimensional organization of the subventricular germinal zone in the adult mammalian brain. J. Neurosci. 17, 5046–5061 (1997).

48. Wittmann, M. et al. Synaptic activity induces dramatic changes in the geometry of the cell nucleus: Interplay between nuclear structure, histone H3 phosphorylation, and nuclear calcium signaling. J. Neurosci. 29, 14687–14700 (2009).

49. Guerrero-Cázares, H. et al. Cytoarchitecture of the lateral ganglionic eminence and rostral extension of the lateral ventricle in the human fetal brain. J. Comp. Neurol. 519, 1165–1180 (2011).

50. Cebrián-Silla, A. et al. Unique organization of the nuclear envelope in the post-natal quiescent neural stem cells. Stem Cell Rep. 9, 203–216 (2017).

51. De Leeuw, R., Gruenbaum, Y. & Medalia, O. Nuclear Lamins: Thin Filaments with Major Functions. Trends Cell Biol. 28, 34–45 (2018).

52. Ochs, R. L., Lischwe, M. A., Spohn, W. H. & Busch, H. Fibrillarin: a new protein of the nucleolus identified by autoimmune sera. Biol. Cell. 54, 123–133 (1985).

53. Waddington, C. H. The Strategy of the Genes. A Discussion of Some Aspects of Theoretical Biology (Allen & Unwin, 1957).

54. Raff, M. Adult stem cell plasticity: fact or artifact? Annu. Rev. Cell Dev. Biol. 19, 1–22 (2003).

55. Rajagopal, J. & Stanger, B. Z. Plasticity in the adult: How should the waddington diagram be applied to regenerating tissues?. Dev.Cell. 36, 133–137 (2016).

56. Kalluri, R. & Weinberg, R. A. The basics of epithelial-mesenchymal transition. J. Clin. Invest. 119, 1420–1428 (2009).

57. Pei, D., Shu, X., Gassama-Diagne, A. &. Thiery, J. P. Mesenchymal–epithelial transition in development and reprogramming. Nat. Cell Biol. 21, 44–53 (2019).

58. Yang, J. et al. EMT International Association (TEMTIA). Guidelines and definitions for research on epithelial-mesenchymal transition. Nat. Rev. Mol. Cell Biol. 21, 341–352 (2020).

59. Reid, A. & Tursun, B. Transdifferentiation: do transition states lie on the path of development? Curr. Opin. Syst. Biol. 11, 18–23 (2018).

60. Alvarez-Dolado, M. et al. Fusion of bone-marrow-derived cells with Purkinje neurons, cardiomyocytes and hepatocytes. Nature. 425, 968–973 (2003).

61. Weimann, J. M., Johansson, C. B., Trejo, A. & Blau, H. M. Stable reprogrammed heterokaryons form spontaneously in Purkinje neurons after bone marrow transplant. Nat. Cell Biol. 5, 959–966 (2003).

62. Magrassi, L. et al. Induction and survival of binucleated Purkinje neurons by selective damage and aging. J. Neurosci. 27, 9885–9892 (2007).

63. Altman, J. Autoradiographic investigation of cell proliferation in the brains of rats and cats. Anat. Rec. 145, 573–591 (1963).

64. Das, G. D. Binucleated neurons in the central nervous system of the laboratory animals. Experientia. 33, 1179–1180 (1977).

65. Ribak, C. E. & Seress, L. Five types of basket cell in the hippocampal dentate gyrus: A combined Golgi and electron microscopic study. J. Neurocytol. 12, 577–597 (1983).

66. Portiansky, E. L., Barbeito, C. G., Flamini, M. A., Gimeno, E. J. & Goya, R. G. Presence of binucleate neurons in the spinal cord ofyoung and senile rats. Acta Neuropathol. 112, 647–649 (2006).

67. Jimenez-Acosta, M. A., Hernandez, L. J. R., Cristerna, M. L. P., Tapia-Ramirez, J. & Meraz-Rios, M. A. Review: Neuronal differentiation protocols of mesenchymal stem cells. Adv. Biosci. Biotechnol. 13, 15–71 (2022).

68. Eguizabal, C., Montserrat, N., Veiga, A. & Izpisua-Belmonte, J. C. Dedifferentiation, transdifferentiation, and reprogramming: future directions in regenerative medicine. Semin Reprod Med. 31, 82–94 (2013).

69. Mollinari, C., Zhao, J., Lupacchini, L., Garaci, E., Merlo, D. & Pei, G. Transdifferentiation: a new promise for neurodegenerative diseases. Cell Death Dis. 2018 Aug 6;9(8):830. doi: 10.1038/s41419-018-0891-4.

70. Kosodo Y. Interkinetic nuclear migration: beyond a hallmark of neurogenesis. Cell Mol. Life Sci. 69, 2727–2738 (2012).

71. Spear, P. C. & Erickson, C. A. Interkinetic nuclear migration: a mysterious process in search of a function. Dev Growth Differ. 54, 306–316 (2012).

72. Hoffmann, K., Sperling, K., Olins, A. L. & Olins, D. E. The granulocyte nucleus and lamin B receptor: Avoiding the ovoid. Chromosoma. 116, 227–235 (2007).

73. Kachi, K. & French, S. W. The connection between the nuclei of binucleated hepatocytes: An ultrastructural study. J. Submicrosc. Cytol. Pathol. 26, 163–172 (1994).

74. Paradis, A. N., Gay, M. S. & Zhang, L. Binucleation of cardiomyocytes: The transition from a proliferative to a terminally differentiated state. Drug Discov. Today. 19, 602–609 (2014).

75. Kanda, T., Sullivan, K. F. & Wahl, G. M. Histone-GFP fusion protein enables sensitive analysis of chromosome dynamics in living mammalian cells. Curr. Biol. 8, 377–385 (1998).

